# Selective binocular integration in mouse visual cortex: binocular but not monocular neurons respond distinctly to congruent stimuli

**DOI:** 10.64898/2026.09.09.750336

**Authors:** Bassler Mathis, Huis In’t Veld Gerjan, Suzuki Mototaka, Pennartz Cyriel

## Abstract

Even though binocular integration is a foremost example of integrative computations in the brain, it remains unknown how the visual cortex mediates it. Here, we investigated neuronal responses to binocular stimuli in layers 2/3 of mouse primary visual cortex (V1) and lateromedial (LM) area with *in vivo* two-photon imaging. We presented mice with three types of binocular stimuli: congruent gratings, orthogonal gratings and a flashing Mondrian pattern in binocular conflict with a grating. We looked for congruency-specific responses to the binocular stimuli, but found that most neurons responded homogeneously across stimulus types. Responses of cells with a strong monocular preference for the contralateral eye responded similarly to monocular and binocular stimulation, while responses of cells with a preference for ipsilateral stimuli showed a strong but nonspecific suppression to all binocular stimuli. Only cells without a strong monocular preference (i.e. binocular cells) exhibited linear integration of monocular responses to incongruent stimuli and binocular facilitation to congruent gratings. Responses to ipsilateral components of a binocular display were found to be poor predictors of binocular responses regardless of eye preference of individual cells. We conclude that most neuronal responses to binocular stimuli in mouse V1 & LM are uninformative about the congruency of the monocular components, with the exception of binocular cells which show either linear or supralinear integration.

**Significance statement:** The combination of signals from both eyes by the visual cortex subserving the genesis of a unified percept poses a main paradigm for studying integrative processes in the brain. A key question in this regard is how binocular stimulus (in-)congruency affects stimulus processing. In this study, we investigated responses to congruent and incongruent signals in mouse primary visual cortex and adjacent lateromedial area with two-photon imaging. We find that while neurons with a strong monocular preference can be affected by binocular stimulus presentation, only neurons without strong monocular preference (i.e. binocular cells) process signals in a congruency-dependent manner.

## Introduction

Binocular integration is a prominent form of integration studied in the field of perception and consciousness science (Pennartz, 2015; Rosenberg et al., 2025). An open question of binocular integration is whether neuronal responses to binocular stimuli in early visual areas depend on the congruency of the monocular images. While initial studies on mammalian binocular processing were predominantly conducted in cats and primates (Hubel & Wiesel, 1962, 1968), the mouse has more recently emerged as a model organism to study binocular processing in early visual areas.

Early studies on binocular processing in early visual cortical areas found notable differences between mammals with frontally facing eyes (such as cats or primates) and mice. One major difference concerns the absence of ocular dominance columns in mouse primary visual cortex (V1) (Dräger, 1975) (but see (Goltstein et al., 2025)). Another major difference concerns the fraction of uncrossed fibers at the optic chiasm, which is very low in mice (<10%) and which, besides callosal projections (Zhao et al., 2013), is a major determinant of binocular processing in V1 (Leinonen & Tanila, 2018). Consequently, contralateral signals dominate even in the binocular part of mouse V1 (Dräger, 1975; Salinas et al., 2017). However, similarities to other mammals have also been found: like in cats, it has been shown that the amount of eye-specific signals in mouse V1 is experience-dependent (Dräger, 1978; Gordon & Stryker, 1996). Furthermore, as in other mammals (Barlow et al., 1967; Poggio & Fischer, 1977), neurons with a sensitivity to binocular disparity have been found in mouse V1 and secondary cortical visual areas, suggesting that mice can extract depth information from binocular signals (Fu et al., 2023; La Chioma et al., 2019, 2020; Scholl et al., 2013).

One question that remains open in mice as well as other mammals is whether binocular response integration depends on congruency of monocular stimuli. An early hypothesis put forward by (Ohzawa & Freeman, 1986) suggested that mammalian V1 cell responses reflect the linear sum of monocular inputs but, over the years, a more nuanced picture of binocular response integration in V1 has emerged (Cox et al., 2019; Fu et al., 2023). In the mouse, studies point towards sublinear integration of congruent stimuli, that is, neurons tend to respond to congruent stimuli less strongly than the sum of responses to the monocular stimuli of the display would suggest (Fu et al., 2023; Longordo et al., 2013; Zhao et al., 2013). Until very recently, binocular responses in mouse V1 were only studied using congruent stimuli, however, in the last year, studies have emerged that also employed binocularly incongruent stimuli (Bassler et al., 2026; Montgomery et al., 2026; Timplalexi et al., 2025). Surprisingly, one of these studies reports that incongruent grating stimuli evoke larger responses in mouse V1 L2/3 pyramidal neurons than either monocular or congruent gratings (Montgomery et al., 2026).

In this study, we investigated whether the responses of mouse V1 and lateromedial area (LM; a primate V2 homolog, (Wang & Burkhalter, 2007)) layer 2/3 (L2/3) cells as recorded with *in vivo* 2-photon microscopy reflect binocular congruency using three types of binocular stimuli: (1) congruent gratings, (2) orthogonal gratings, and (3) a grating in binocular conflict with a changing (“flashing”) pattern of rectangles (“Mondrian”). The first two conditions are classical stimuli for studying orientation-tuned responses, while the last stimulus is inspired by the human paradigm of Continuous Flash Suppression (Tsuchiya & Koch, 2005). We find that most L2/3 cells in V1 and LM respond homogeneously to binocular stimuli regardless of congruency of the monocular components. Responses of cells with a strong preference for the contralateral eye responded similarly to monocular and binocular stimuli while those preferentially responding to the ipsilateral eye were strongly suppressed across all binocular conditions. Only the group of binocularly responsive neurons exhibited near-linear integration of monocular signals, which in the case of congruent stimulation was even supralinear.

## Materials and Methods

### Animals

All animal experiments were performed according to the national and institutional regulations. Our experimental protocol was approved by the Dutch Commission for Animal Experiments (CCD application number: AVD11100202216078) and by the Animal Welfare Body of the University of Amsterdam. We used 5 (1 female) double transgenic mice (background: C57BL/6J) from in-house breeding of Rasgrf2-2A-dCre (JAX #022864, Cre-driver line) and Ai148D (JAX #030328, GCaMP6f reporter liner) (Daigle et al., 2018; Song et al., 2017). Both lines were originally purchased from the Jackson Laboratory (Bar Harbor, Maine, USA). Mice were between 8 and 52 weeks old during experiments. They were housed socially on a reversed day night cycle (lights on: 8 pm, lights off: 8 am) and had free access to water and food. GCaMP6f expression was induced by injecting mice with 300 mg / kg trimethoprim (TMP; T883-5G, Sigma Aldrich) dissolved in dimethylsulfoxide (DMSO) intraperitoneally (IP, concentration: 250 mg / ml) once per day over a series of three days.

### Surgery

For two-photon imaging, mice were surgically implanted with a circular 4-mm cranial window over the left posterior hemisphere as well as a custom-built titanium head bar for head fixation. At the start of the surgery, anesthesia was induced with 4-5% isoflurane in 100% oxygen and a constant supply of 1-2% isoflurane was maintained throughout the surgery. Additionally, mice were subcutaneously injected with 10 mg / kg carprofen (Carprofelican, Decan) for analgesic purposes. Mouse body temperature was maintained with a heating pad, temperature probe and controller (TCAT-2LV, Physitemp instruments). After induction, the anesthesia depth was assessed by testing the pedal withdrawal reflex. Then, the fur on the head of the mouse was shaved off, and an antiseptic (Betadine, Mylan) and a topical anesthetic (Xylocaine, Aspen) were applied before removing part of the skin above the skull with scissors and forceps. The head bar contained a circular opening which was positioned above the target location of the cranial window. The head bar was first provisionally glued to the skull (Pattex Sekundenkleber, Henkel), then firmly attached with dental cement (C&B Superbond, Sun Medical). A dental drill was used to create a circular craniotomy slightly exceeding 4 mm in diameter on the left posterior skull such that the posterior part of the craniotomy barely exceeded the lambdoid suture. We were especially careful while drilling over the suture as a large blood vessel, the transverse sinus, lies beneath it which must not be punctured. The skull piece within the craniotomy was removed and the exposed cortex was flushed with saline. The cranial window was placed inside the craniotomy, then fixed with glue and dental cement (Simplex Rapid, Kemdent). The exposed skull was covered with more dental cement and a small metal ring (diameter: 1.8 cm, height: 3mm) was fixed to the head bar for light shielding during imaging. Mice were provided with an analgesic (carprofen) and antibiotics (enrofloxacin, Baytril, Bayer) for three days after surgery by mixing the components into the drinking water.

### Data collection Binocular setup

The binocular setup used in this study was first described in (Bassler et al., 2026). Briefly, two projectors (ML1050ST, Optoma) were mounted on a custom table and pointed at a polarization-preserving screen (ST-Pro-X, Screen Tech-Shop). A linear polarization filter (XP42-18, Edmund Optics) was positioned in front of each projector lens as well as each eye of a mouse. The polarization filters in front of the projectors were held by custom 3D-printed holders while the ones in front of the mouse’s eyes were held by custom 3D-printed mouse glasses attached to a stereotactic arm (LC mini holder LC5200, Noga). The polarization axes of the filters were carefully aligned such that light from one projector could only reach one eye. Alignment was verified using cameras and a custom-built photometer.

### Microscope

Two-photon and widefield recordings were performed using a Leica DM6000B microscope. For two-photon recordings, a Spectra-Physics Mai Tai mode-locked Ti:sapphire laser with a wavelength of 920 nm was used and a plane of 570 x 570 μm at a depth of 100 – 200 μm was imaged through a 16x objective (Leica HC FLUOTAR L 16x/0.60 IMM CORR VISIR) at a frequency of 27.5 Hz (bidirectional scanning). For previous studies using the same setup, see (Bassler et al., 2026; Goltstein et al., 2015; Montijn et al., 2016). For widefield imaging, we recorded the brain surface under green LED light at 470 nm (Thorlabs M470L5 LED with Thorlabs T-Cube driver) through the cranial window using a 1.25x objective (Leica HC PL FLUOTAR 1.25x/0.04 T), filter cube (Leica I3 DM 513828) and camera (Basler ace 2 a21920-60um).

### Retinotopic mapping

Before starting two-photon sessions, we presented anesthetized mice with moving bar stimuli while recording the full cranial window with widefield imaging to chart the visual areas inside the window. Mice were anesthetized with isoflurane (see above) and additionally received an injection of 5 mg / kg xylazine S.C. to prevent eye drift (Nair et al., 2011). As xylazine lowers the amount of isoflurane needed for anesthesia, and as we were aiming for light anesthesia to record neural signals, isoflurane levels were kept low, between 0.5 and 1%. Mouse body temperature was controlled during imaging the same way as during surgeries.

For stimulus presentation, we adopted the retinotopic mapping paradigm described in (Zhuang et al., 2017). Briefly, flickering checkerboard bars (width: 20 visual degrees, square size: 25 visual degrees) moved across a large computer screen (with a grey background) presented to the right eye. Bars could move in one of four directions (left to right, right to left, bottom to top and top to bottom). Each bar sweep was repeated 20 times. Bar sweeps were interspersed with intertrial intervals of 2-3s (grey background only). The analysis of the widefield imaging data obtained during retinotopic mapping stimulation is described in the section “Identification of visual areas”.

### Binocular visual stimulation

All stimuli were generated using custom Python 3 code and the PsychoPy package (Peirce et al., 2019) and shown on the aforementioned binocular setup. Mice were presented with 1s stimuli on a grey background with interspersed intertrial intervals of 1.5 to 2.5s (grey background only).

Monocular stimuli consisted either of circular sinusoidal drifting gratings (30 visual degrees size, 0.1 cpd spatial frequency, 2 Hz temporal frequency) or of a group of pseudorandomly arranged and sized “rectangles with pseudorandomly assigned greyscale intensities (35 visual degrees total size, 5 to 15 visual degree size per rectangle, overlay of 50 rectangles). Monocular stimuli could be presented either to the left or right eye using our binocular setup. Stimuli shown via either projector were carefully aligned such that they covered the same extent of visual space. We refer to the rectangle display as a Mondrian following Continuous Flash Suppression literature (Tsuchiya & Koch, 2005). The arrangement of rectangles that made up the Mondrian was changed (“flashed”) every 100 ms which is below the critical flicker frequency of mice (Umino et al., 2018). The sequence of rectangle configurations was generated once at the beginning of each recording session and then repeated throughout the session. For gratings, four drift directions were possible (0, 90, 180 or 270°).

Grating stimuli could be presented to one eye only (monocular grating = MG condition) or on both eyes (congruent gratings = CG or orthogonal gratings = OG condition). In the CG condition, both gratings had the same drift direction (and were thus fully congruent as the name suggests). In the OG condition, gratings had an interocular drift direction difference of 90°. If the screen was viewed without polarization filter glasses, the CG stimulus appeared as a single grating while the OG stimulus appeared as a plaid (Fig. S1). Furthermore, a Mondrian stimulus could be shown alone (monocular Mondrian = MM condition) or in binocular conflict with a grating (CFS condition).

There were 26 stimulus displays in total: eight MG variants (two eyes x four drift directions), two MM variants (left or right eye), four CG variants (four drift directions), four OG directions (0° x 90°, 90° x 0°, 180° x 270°, 270° x 180°; right x left eye) and eight CFS variants (two eyes x four drift directions). For a visualization of all stimulus displays, see Fig. S2. . The stimuli were shown in blocks of repeats (20 repeats per stimulus) where each block contained the 26 stimulus displays in a new pseudorandom order. Stimuli were presented at the horizontal center of the visual field of the mouse at an elevation of 10 visual degrees.

### Two-photon imaging

Mice were first habituated to sitting in a cylindrical tube while being head-fixed before starting imaging sessions. This habituation started with very short periods of head fixation per day (1 – 2 min) after which head fixation duration was increased over several days. The mouse head bar contained two screw holes which could be lined up with corresponding screw holes on a custom head bar holder such that the head bar could be fixed to the holder with screws. Once mice were comfortable with being head fixed for a prolonged period, they were further habituated to having the mouse glasses positioned close to their eyes.

Before an imaging session, the cranial window was cleaned carefully with distilled water, then covered with a mixture of ultrasound gel and water. The head bar holder with the mouse was placed on the microscope stage, after which the mouse glasses were positioned in front of the eyes of the mouse. A custom 3D printed shielding tube was attached to the microscope objective, then the objective was positioned over the cranial window. The shielding tube was connected to the implanted shielding ring to avoid interference of stimulus light with two-photon recordings. The binocular setup was positioned in front of the microscope and the center of its screen was aligned to the horizontal center of the mouse’s visual field. We then started stimulus presentation and two-photon imaging. Throughout imaging, we monitored the behavioral state of the mouse with two cameras.

### Analysis

All data were analyzed using custom code in Python 3. We mention relevant packages where they apply.

### Multiple comparison correction

Whenever multiple statistical tests were conducted (except for Tukey HSD tests which have inherent multiple comparison correction), p-values were corrected for multiple comparisons by Benjamini-Hochberg correction using the *pingouin* package unless specified otherwise.

### Identification of visual areas

We analyzed the widefield video data obtained from retinotopic mapping as described in (Zhuang et al., 2017). Briefly, we averaged widefield calcium imaging movies of bar sweeps of the same direction, then computed the Fourier transform (*scipy.ffi.rffi*) at the bar sweep frequency to obtain a phase map for each direction. The phase maps revealed to which position of the bar each part of the cortical surface responded to. We subtracted phase maps of the same bar orientation (but opposing movement direction) from each other to obtain azimuth and altitude maps. Finally, the azimuth and altitude maps were combined to form a visual field sign map which depicts areas of shared retinotopy. We identified brain areas V1 and LM based on the visual field sign maps and comparison to the reference map given by (Zhuang et al., 2017).

### Cell extraction

Cell regions-of-interest (ROIs) were extracted from two-photon videos using Suite2p (Pachitariu et al., 2017). Before using Suite2p, the two-photon videos were motion-corrected using the NormCorre algorithm from the CaImAn package (Giovannucci et al., 2019). Then, a custom algorithm was employed to remove bidirectional scanning artefacts. We used Suite2p’s internal cell classifier as well as ROI responses to identify stimulus-selective cells (see section “Cell selection”).

### Signal scaling

Suite2p fluorescence traces were scaled before further analysis. Following (Chen et al., 2013), neuropil activity (F_neuropil_) was first subtracted from ROI fluorescence values (F_0_) as follows:

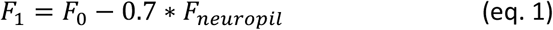

Then, ROI fluorescence values (F_1_) were transformed into dF/F values (F_final_) by scaling the response of each ROI in each trial with the average response during 1s before stimulus onset (baseline period, F_baseline_).

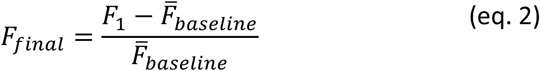

### Cell selection

We filtered ROIs according to four criteria to obtain our selection of grating-selective cells.

1. Somatic activity: The probability of an ROI being the soma of a cell had to exceed 50% as determined by Suite2p’s internal cell classifier. This ensured that ROIs captured cell somata instead of e.g. dendrites.
2. Selective responses to monocular gratings: For each cell, an ANOVA was conducted with the *scipy* package using the responses to eight monocular gratings as groups with twenty stimulus repetitions per group. Each data point was the average response during 1s stimulus presentation of a trial. The ANOVA tested whether varying monocular grating type (eye of presentation and drift direction) modulated the response strength more than the repetition of varying monocular grating repetitions. A positive outcome (p < 0.05) indicated that a cell responded selectively to the monocular gratings. As the stimulus repetitions were distributed evenly over the duration of a recording, this test also ensured temporal stability of responses. ANOVA p-values were corrected for m = 10,680 ROIs. For a similar approach to assessing grating selectively, see (Chen et al., 2013).
3. Strong responses to preferred gratings: The average response (averaged over 1s stimulus duration and 20 repetitions) to the preferred grating of a cell (which was the monocular grating that evoked the highest average response) had to exceed the arbitrary threshold of 0.25 dF/F. This ensured that cells responded strongly to their preferred grating.
4. Low baseline noise: The variance during 1s before the stimulus period across all trials had to be lower than 1 dF/F. This removed ROIs with very noise fluorescence traces such as those with very low baseline fluorescence, which could result in unstable dF/F values.

### Decoding

#### Stimulus condition decoding

We trained linear support vector machines (SVMs) to decode stimulus condition from the average response in each trial during 1s stimulation using the *scikit-learn* package. We arranged the average responses of each cell during 1s of trials of the MG, CG, OG and CFS type as a vector, using only trials that included a cell’s preferred monocular grating in the visual display (80 trials per cell). We z-scored the responses in that vector, then trained an SVM for each of the four stimulus conditions to predict whether a trial came from that stimulus condition (binary classification with 10-fold cross-validation). Decoding performance was measured as decoding accuracy and significance of decoding performance was assessed by permutation testing using 200 shuffles per fitted SVM, with a cell classified as part of the decodable fraction when p < 0.05. No multiple comparison correction was used for the permutation tests.

#### Statistical assessment of coding fraction differences

To compare coding fractions across conditions (Fig. 1H), we bootstrapped the fractions of cells from which a stimulus condition could be decoded above chance using 10,000 bootstrap samples. For each bootstrap iteration, we computed the pairwise difference Δ in coding fractions f_i_ and f_j_ (where i and j denote two separate stimulus conditions) as follows:

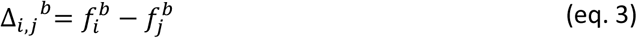

where b indicates the bootstrap iteration. Then, we assessed the fraction of bootstrap iterations for which that Δ was positive (> 0) or negative (=< 0).

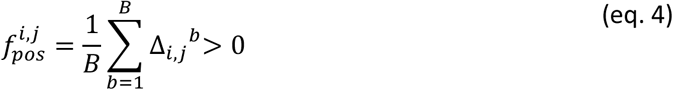

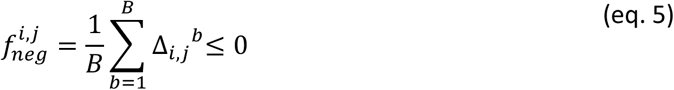

**Figure 1.**
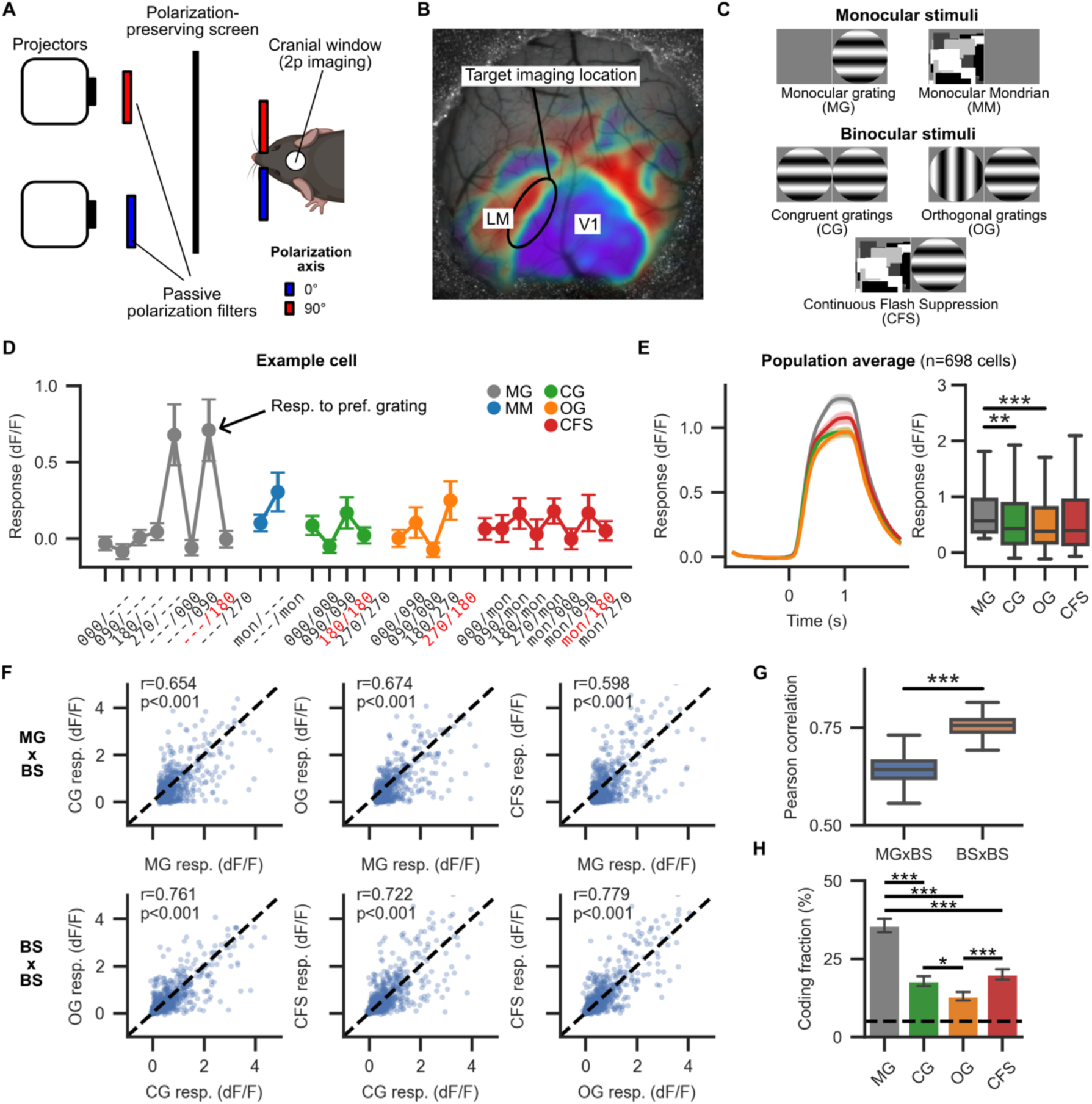
Neurons in mouse primary visual cortex and lateromedial area show net suppression of visual responses by binocular stimuli regardless of stimulus congruency. (A) Mouse binocular stimulation setup based on linear polarization filters. (B) Target two-photon imaging region at the border of V1 and LM drawn over a retinotopic map of one of the mice used in the study. The retinotopic map depicts the visual field sign (blue, red; regions of unchanging field sign values have a shared retinotopy and thus form a visual area) and is overlaid on an anatomical widefield image of the cranial window (grey). For details of the retinotopic mapping, see Methods. (C) Depiction of stimulus conditions. We showed mice two types of monocular stimuli (a grating or a Mondrian) and three types of binocular stimuli (congruent gratings, orthogonal gratings, continuous flash suppression). Stimulus duration was 1s followed by an intertrial interval of 1.5 to 2.5s. Grating stimuli were slowly drifting, sinusoidal gratings with one of four drifting directions (0, 90, 180 or 270°). Mondrian stimuli consisted of pseudorandom grayscale patches whose arrangement was changed (“flashed”) every 100 ms, both during monocular and binocular presentations. Each stimulus condition was repeated 20 times. For a full overview of all 26 conditions, see Fig. S2. (D) Average responses of an example cell to every stimulus condition during 1s of stimulus presentation. Error bars indicate SEM. The stimulus conditions are encoded as “right eye stimulus/left eye stimulus”. A three-digit number indicates that a grating stimulus was displayed with that orientation, “mon” indicates that a Mondrian stimulus was displayed and “---” indicates the grey background was displayed. X-tick labels of stimulus conditions that include this cell’s preferred grating are highlighted in red. (E) Population response of 698 grating-selective V1 and LM cells to four stimulus types. On average, cell responses to binocular stimuli were suppressed compared to the monocular condition. There were no significant differences in response amplitudes between different binocular conditions. Because neurons in mouse early visual areas tend to be strongly orientation-tuned, here and in the following, for each cell, only stimulus conditions that included a cell’s preferred monocular grating were included in the analysis. Left: average responses over time. Right: Boxplot of responses. Time = 0 s indicates stimulus onset. Shaded regions around the means indicate the SEM. ***: p < 0.001, **: p < 0.01, Tukey HSD after ANOVA. (F) Scatterplots and Pearson correlation coefficients (r) between responses to monocular and binocular conditions, with their p-values. Each dot represents the average dF/F response of a cell in a condition. MG x BS: correlation of monocular grating responses with one of three binocular stimulus condition responses. BS x BS: correlation of two binocular stimulus condition responses. (G) Cell responses to binocular stimuli of different conditions were more strongly correlated with each other than with responses to monocular stimuli. ***:p < 0.001, bootstrap hypothesis test, 1000 shuffles. (H) Fraction of cells from whose responses the stimulus condition rendered on the abscissa was significantly decodable (i.e., above chance level) as assessed by permutation testing (200 permutations per cell). No multiple comparison correction was used for permutation testing. Cells were most informative about whether the stimulus display was monocular or binocular, but less informative about the type of binocular display. The dashed line indicates chance level (5%). Error bars indicate bootstrapped standard deviation (10,000 bootstraps). **: p < 0.05, ***: p < 0.001, bootstrap hypothesis test with MCC.

Here, B indicates the total number of bootstraps (10,000).

Finally, we determined the significance of a fractional difference with the following p-value:

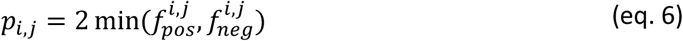

For Fig. 1H, this procedure resulted in six p-values (pairwise comparison of four conditions). For Fig. 2G, we repeated the above procedure but now also split cells into eye-preference groups, resulting in 12 coding fractions (four conditions, three eye-preference groups) and 66 pairwise comparisons. In both cases, the p-values were adjusted for multiple comparisons as described above.

**Figure 2.**
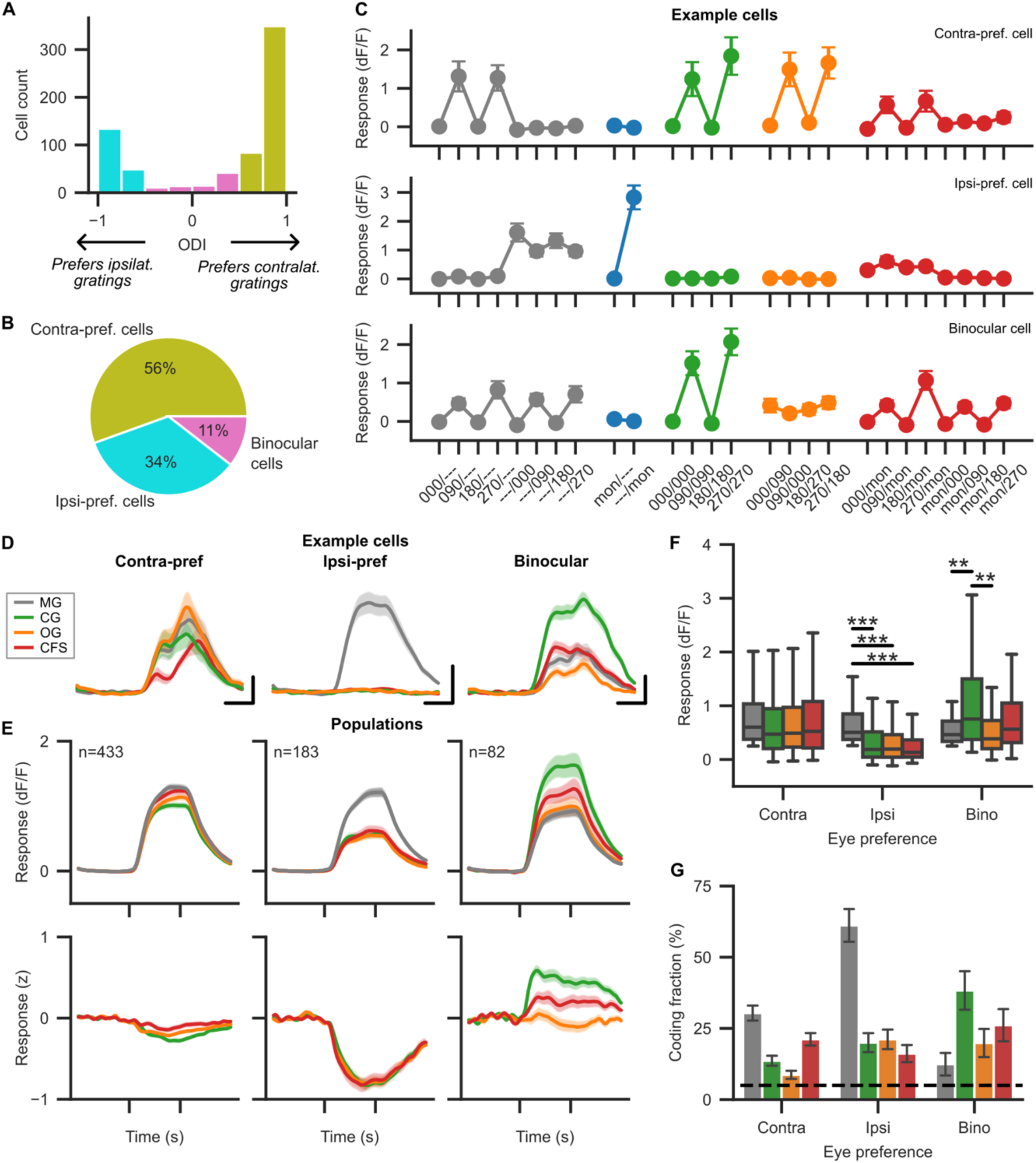
Only binocularly responsive cells respond selectively to congruency differences. (A) We divided cells into contralateral-preference, ipsilateral-preference and binocular cells based on ocular dominance index. The average ODI was 0.33 ±0.03 (mean ± SEM). (B) Only a small minority of cells did not have a strong monocular preference (0.5 > ODI > 0.5). We call these cells binocular cells. (C) Responses of three example cells with different eye preferences to all stimulus conditions. Convention as in Fig 1D. Only the binocular cell shows selective facilitation by congruent stimuli. (D) Average responses of the same example cells as in C but now only including stimulus presentations that included each cell’s preferred gratings. The horizontal scale bar indicates 0.5s, the vertical scale bar indicates 1 dF/F. (E) Responses of eye-preference populations. Conventions as in Fig 1E. In the lower row, before calculating the population average, for each cell and binocular condition, the average MG response was subtracted from the binocular response and the result was scaled by the pooled standard deviation of the MG and binocular response. (F) Differences in response amplitudes per eye preference group. . **: p < 0.01, ***: p < 0.001, Tukey HSD test. To avoid overcrowding the panel, significant differences in response strengths between eye preference groups are omitted (G) Fractions of cells from which an individual stimulus condition is decodable above chance per eye-preference population. Contra: contra-preferring cells, Ipsi: ipsi-preferring cells, Bino: binocular cells. Conventions as in Fig 1H.

#### Brain area decoding

For Fig. S2, we tested whether the brain area label of all cells could be decoded based on their activity patterns in the 26 stimulus conditions. The average responses during 1s of stimulation across all twenty repeats per condition were used as features, resulting in a matrix of shape [698 cells, 26 average responses]. We again used a linear SVM with 10-fold cross-validation, and assessed decoding performance by permutation testing with 100 permutations. The brain area label decoding performance was found to be at chance level, indicating that brain area information was not contained in the cell responses.

### Linear regression

For Figures 3 and 4, we fit linear regression models using the *statsmodels* package. Regression models always had only one predictor variable (Fig 3.: linear sum of monocular responses, Fig. 4: either contralateral or ipsilateral stimulus response) and an offset. Both the dependent and independent variables were vectors where each element consisted of the average response of a cell (as averaged over 1s stimulation and 20 repetitions per stimulus condition). For Fig. 3, we employed one-sample t-tests to convert slope estimates into t-values. Specifically, we determined the t-value by t = (slope - 1) / SE, where SE is the standard error of the slope and 1 indicates the null hypothesis that linear sums predicted binocular responses with a weight of 1. We then converted the t-value into a p-value using the t-distribution. For Fig. 4, we obtained bootstrapped coefficients of determination (r^2^) for each fit with 1000 bootstrap samples. To compare r^2^ values, we used bootstrap hypothesis testing similar to the approach described for comparing coding fractions. Briefly, we computed the difference in bootstrapped r^2^ values between contra- and ipsilateral stimulus response predictors per bootstrap iteration (abbreviated: CSR, ISR). We determined the fraction of bootstrapped differences where r^2^_CSR_ exceeded r^2^_ISR_ and vice versa. We then converted these fractions into a p-value using: p = 2 * min(fraction_CSR>ISR_, fraction_ISR>CSR_).

**Figure 3.**
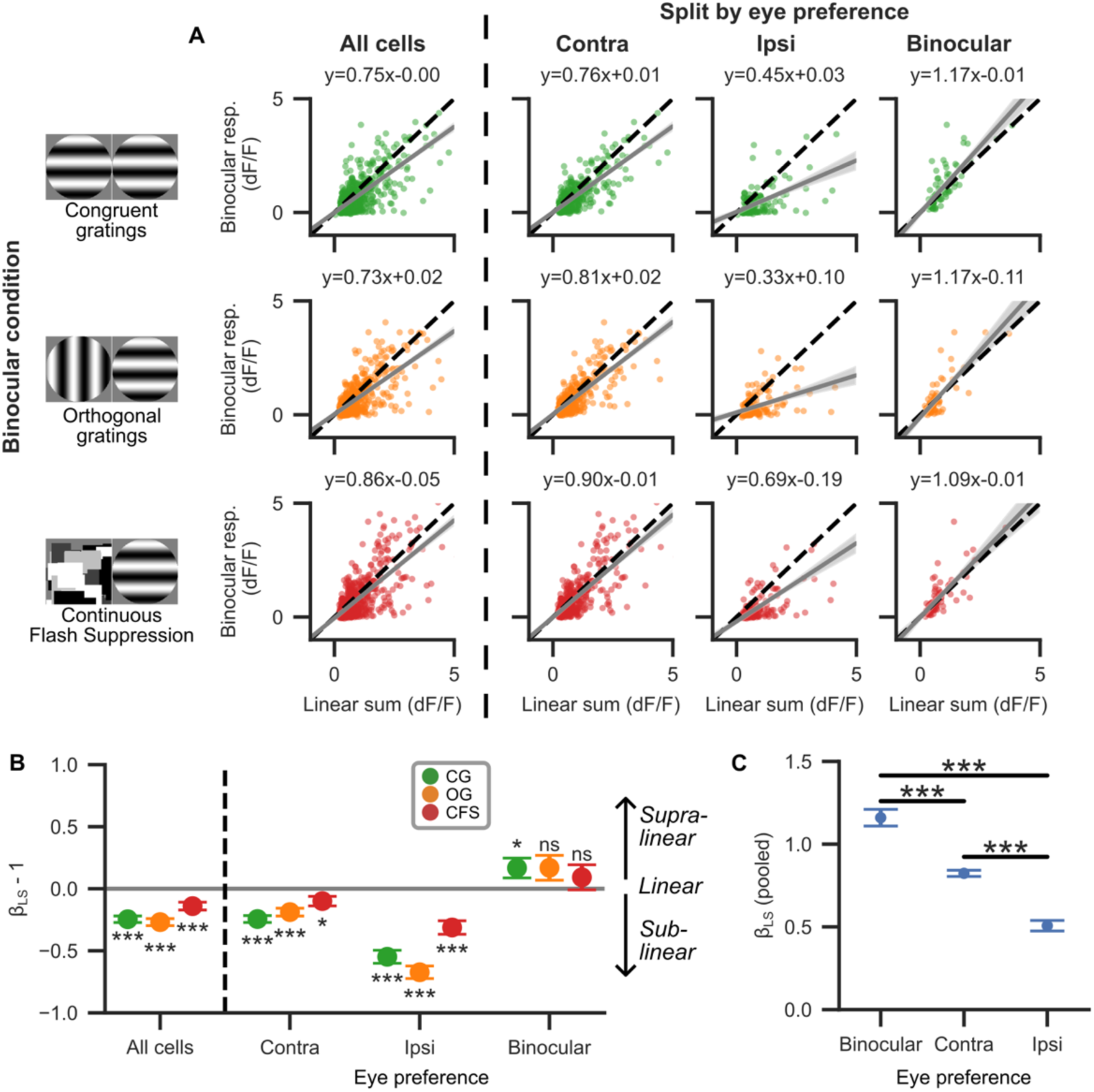
Cells with strong monocular preference integrate sublinearly, binocular cells linearly or supralinearly. (A) Comparison of linear sums of responses to monocular stimuli of binocular displays with actual binocular responses. Grey lines are linear regression lines to predict binocular responses from the sum of monocular responses. The best-fit regression formula is given for each subplot as y = β_LS_ * x + β_offset_, where y is the binocular response and x is the linear sum of the monocular component responses. Each dot represents the average activity of a cell. Shaded areas around the linear regression indicate the 95% confidence interval. Black dashed line indicates 1:1 linearity (β_LS_ =1). (B) Cell populations with strong monocular preference (contra- or ipsilateral) show sublinear integration, while those without strong preference (i.e. binocular cells) show either linear or supralinear integration. Error bars indicate the standard deviation of the slope estimates. ***: p < 0.001, **: < 0.01, *: p < 0.05, ns: p >= 0.05, one-sample t-test against β_LS_ =1, MCC for 12 tests. (C) When pooling binocular responses across conditions and rerunning the regression analysis from (A), differential signal integration by the eye preference groups is revealed. All response integration strengths as measured by slope β_LS_ differ significantly (p < 0.001, two-sided t-test for all three pairwise comparisons, MCC for three tests).

**Figure 4.**
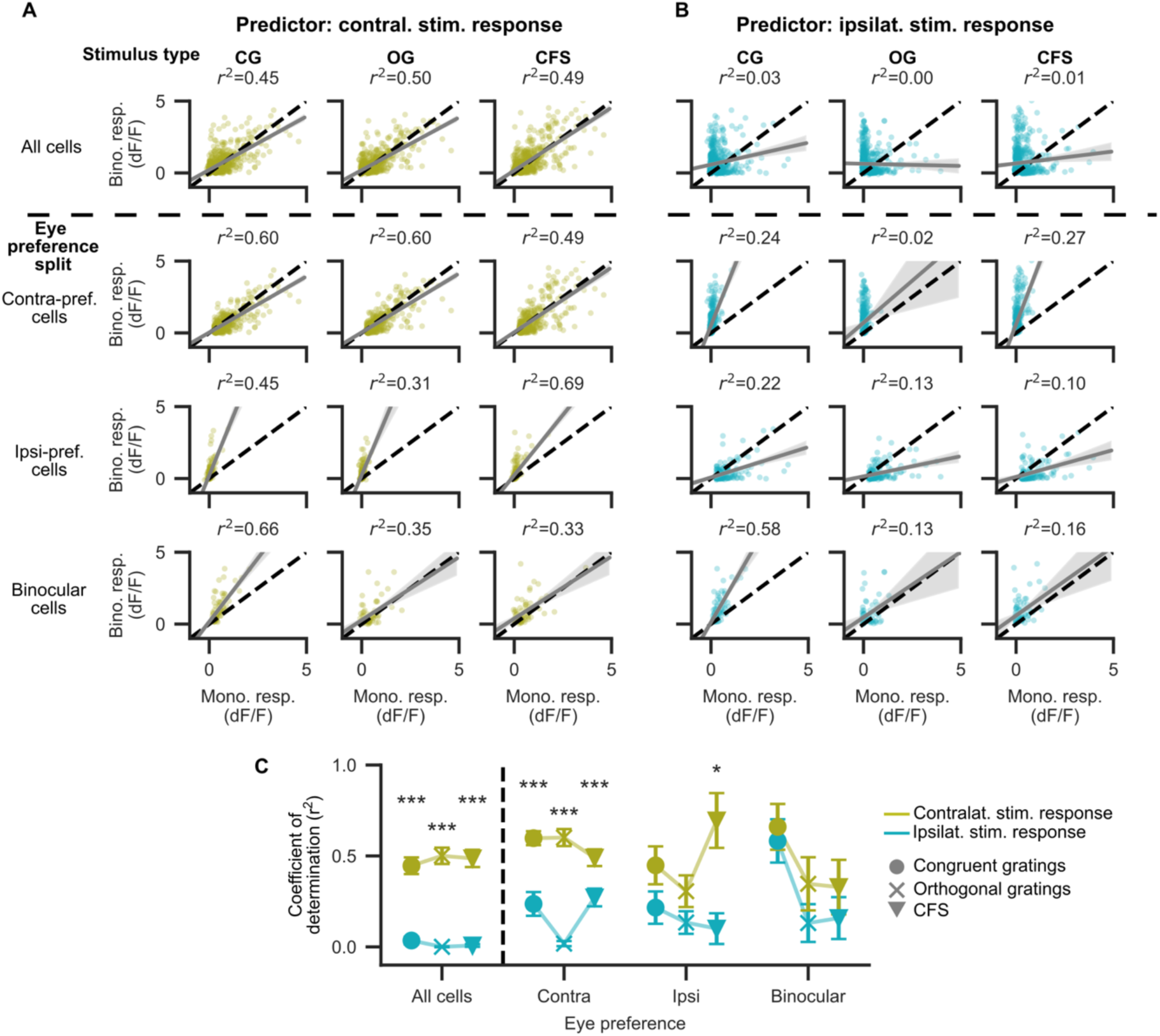
Responses to ipsilateral, but not contralateral stimuli are poor predictors of binocular responses. (A) Linear regression of binocular response (y-axis) and response to contralateral stimulus (x-axis). Grey lines are linear regression lines, the grey shaded region indicates the 95% confidence interval. Note that confidence intervals may become large for regions with few data points. Each dot is one cell. In these linear regressions, the coefficient of determination r^2^ values atop each plot measures how well binocular responses can be predicted from monocular ones. The dashed black line indicates linear scaling between contralateral and binocular responses. Mono. / Bino. resp.: monocular / binocular response. (B) Same as (A) but now using ipsilateral stimulus responses as predictors for binocular responses. (C) Statistical comparison between r^2^ values of the contralateral and ipsilateral stimulus responses as predictors in the linear regressions from (A). Error bars indicate SD and were estimated with bootstrapping (1000 samples). *: p < 0.05, ***: p < 0.001, bootstrap hypothesis test with 1000 bootstraps between r^2^ values using contra- or ipsilateral stimulus responses as the predictor variable (null hypothesis: r^2^_CSR_ = r^2^_ISR_), MCC for 12 tests.

## Results

To present mice with monocular and binocular stimuli at the same location in visual space, we used a polarization-filter based binocular setup which allowed guiding light from a projector to only one of the two eyes of a mouse with minimal cross-talk (Fig 1A, see also (Bassler et al., 2026)). During stimulus presentation, we imaged the activity of cortical L2/3 neurons at the border of V1 and LM with *in vivo* two-photon imaging in five passively observing mice (Fig 1B). We presented mice with two monocular stimulus types and three binocular stimulus types (Fig 1C). Monocular stimuli consisted either of a slowly drifting sinusoidal grating with one of four drifting directions (“monocular grating”, abbreviated: MG; 0, 90, 180 or 270°, size: 30 visual degrees,) or of a Mondrian stimulus that was made up of pseudo-randomly arranged grayscale rectangles whose arrangement was changed (“flashed”) every 100 ms (size: 35 visual degrees, “monocular Mondrian”, abbreviated: MM). The latter stimulus was inspired by the human paradigm of Continuous Flash Suppression (Tsuchiya & Koch, 2005) and was used by us in a previous study (Bassler et al., 2026). Monocular stimuli could be shown to either the right or left eye while the other eye was presented with a grey background.

Our binocular stimuli differed in binocular congruency, that is, in the amount of similarity between the two images which were presented to the two eyes. Congruent grating (CG) stimuli consisted of the same grating shown to either eye. Orthogonal grating (OG) stimuli still consisted of two gratings but now with an interocular 90° orientation difference, which resulted in incongruency both in the drifting direction and the overlap of black and white stripes. The Continuous Flash Suppression (CFS) condition consisted of a grating on one eye and a flashing Mondrian pattern on the other eye. Since the Mondrian pattern was changed every 100 ms, this stimulation resulted in variable congruency at each subspace of the visual field (Fig. S1). All stimuli were presented for 1s at the same location in visual space over a grey background and interspersed with intertrial intervals consisting of the grey background shown for 1.5 to 2.5s. In total, 26 stimulus conditions were shown, and each condition was repeated 20 times per recording session (Fig. S2). The response of an example neuron to all 26 stimulus conditions is shown in Fig 1D.

From 24 recording sessions, we extracted 10,680 neuronal ROIs with Suite2p (Pachitariu et al., 2017). We filtered ROIs by noise levels and Suite2p cell classification and then looked for cells with differential responses to the eight monocular grating conditions as determined by ANOVA. We call such cells grating-selective. For each grating-selective cell, we determined the monocular grating that evoked the highest response (the preferred grating) and only included cells in our analysis whose average response to the preferred grating exceeded the arbitrary threshold of 0.25 dF/F. The preferred grating was distinguished from other gratings by its eye-of-origin and drifting direction. For all analyses, to avoid averaging over conditions with weak or absent responses, we only included cell responses to stimulation conditions that included each cell’s preferred grating either as the sole stimulus or as one of the two components of a binocular display. Overall, we found 698 grating-selective cells from V1 and LM. Since imaging regions in V1 and LM were close to the interarea-border and we could not distinguish between neurons from both areas with linear classification based on their response patterns (Fig. S3), we pooled the cells from V1 and LM for the remainder of the study.

### Majority of V1/LM cells responds homogeneously to binocular displays regardless of congruency

We first investigated how neurons responded to our stimuli by computing the average response of each neuron to each condition. We found that in the overall population of cells, MG stimuli elicited on average the strongest response, while all binocular condition responses (CG, OG and CFS) were comparatively weaker (Fig. 1E). This effect was significant for the MG-CG comparison (ANOVA p < 0.001, n = 698 cells, Tukey’s HSD p = 0.005) and MG-OG comparison (p < 0.001) and a similar (non-significant) trend was seen for the MG-CFS comparison (p = 0.117), indicating that adding a second stimulus to the binocular display generally had a suppressive effect on grating-selective cells.

Interestingly, despite differences in binocular congruency, we found no significant differences in response amplitudes between the three types of binocular displays (all p > 0.05). When correlating response amplitudes of the four conditions, we found that while all condition pairs showed strong correlations (Pearson correlation coefficient r >= 0.6, p < 0.001, multiple comparison correction [MCC] for 6 tests; Fig. 1F), responses to binocular conditions were more strongly correlated with each other than with responses to monocular gratings (p < 0.001, bootstrap hypothesis test with 1000 shuffles; Fig. 1G). This suggests a tendency for cells to respond homogeneously to binocular displays *regardless of congruency*. To test how informative cell responses were about the stimulus displays, we attempted to decode stimulus types (MG, CG, OG or CFS) from trial responses for each cell. We found that we could decode whether a response was evoked by MG trials from 36% of all cells above chance level (249/689 cells), while binocular conditions could be decoded only from smaller fractions (CG: 18%, OG: 13%, CFS: 20%; Fig. 1H), indicating that the responses of most cells were not informative about binocular stimulus type.

### Only neurons with binocular preference show facilitation to congruent gratings on both eyes

Dominance of contralateral eye signals in the binocular region of mouse V1 is well known (Dräger, 1975; Salinas et al., 2017). Our previous study on binocular conflict processing in V1 and LM showed differential processing depending on whether cells responded preferably to stimuli on the contra- or ipsilateral eye (Bassler et al., 2026). We therefore computed the ocular dominance index (ODI) for all neurons (Fig. 2A) on MG responses to see if we could find congruency-dependent responses in groups of cells with a particular eye preference. An ODI of 1 indicates that a cell responds exclusively to a monocular grating with the preferred orientation on the contralateral eye (with an ODI of -1 indicating the same but for the ipsilateral eye). We divided cells into three eye preference groups: (1) contra-preferring cells, having an ODI larger than 0.5, (2) ipsi-preferring cells, with an ODI smaller than -0.5 and (3) binocular cells, with an intermediate ODI (0.5 > ODI > -0.5). In accordance with previous studies, we found a majority of contra-preferring cells (433/698 cells, 56%). Ipsi-preferring cells made up a smaller fraction of the total population (265/698 cells, 34%), and only a small fraction of cells were binocular (82/698 cells, 11%; Fig. 2B). Responses of three example cells from the three eye preference groups are shown in Fig. 2C and D.

When examining average responses of these different eye-preference groups (two-factor ANOVA with factors stimulus condition and eye preference group, p < 0.001; subsequent Tukey HSD test for 66 pairs), we observed clear differences between the contra- and ipsi-preferring populations. The contra-preferring population responded to all stimulus conditions equally (Fig. 2E and F; p_tukey_ > 0.05 for all condition pairs), while the ipsi-preferring population exhibited suppressed responses to all binocular conditions as compared to the MG condition (p_tukey_ < 0.001 for MG-CG, MG-OG and MG-CFS comparisons). While the responses of the ipsi-preferring population were clearly modulated by binocular stimulation as compared to monocular stimulation, there were no significant differences in response strengths between the different binocular conditions (all p_tukey_ > 0.05 for CG-OG, CG-CFS and OG-CFS comparisons), indicating that the observed suppression was indifferent to congruency. Consequently, while MG responses of contra- and ipsi-preferring responses did not differ (p_tukey_ = 0.890), CG, OG and CFS responses were lower in the ipsi-than in the contra-preferring population (p_tukey_ < 0.001). Notably, congruency-specific responses were only apparent for the binocular cells, for which CG responses significantly exceeded MG and OG responses (Fig. 2F; CG vs MG: p_tukey_ = 0.001, CG vs OG: p_tukey_ = 0.008, CG vs CFS: p_tukey_ = 0.429).

As expected by the condition responses, when trying to decode stimulus type (as in Fig. 1H), responses of contra-preferring cells were largely indiscriminate about the type of stimulus shown (MG: this condition was decodable above chance in 30% of the cells, CG: 13%, OG: 8%, CFS: 21%; Fig 2G). From most ipsi-preferring cells (110/183 cells, 60%), we were able to decode whether stimuli were monocular or binocular (likely because binocular stimulation evoked suppression across all binocular displays). However, the type of binocular stimulus could again be decoded from only a small fraction of ipsi-preferring cells (CG: 20%, OG: 21%, CFS: 16%). For the binocular cells, it was possible to decode congruent stimulation from 38% cells, more than any other condition in that eye preference group (MG: 12%, OG: 20%, CFS: 26%).

### Sublinear response integration is most prevalent in cells with strong monocular preference

Previous studies have reported sublinear binocular response integration in mouse V1 (Longordo et al., 2013; Zhao et al., 2013), but they did not examine different V1 subpopulations based on eye preference. We sought to refine this picture with our eye-preference split as we find that at least binocular cells can exhibit binocular facilitation, which suggests supralinear or linear instead of sublinear integration. We fit linear regression models to the sums of the responses to the monocular stimuli of each binocular display (linear sums or LS in the following) to predict binocular responses (Fig 3A). We took the estimate of the slope of those regression models to indicate whether responses were on average sublinear (slope β_LS_ < 1), linear (β_LS_ = 1) or supralinear (β_LS_ > 1).

As expected from previous studies, we found best fits with slopes below 1 for cells with strong monocular preference across all binocular conditions (p < 0.05, one-sample t-test against β_LS_ = 1, MCC for m = 12 tests, Fig. 3B), indicating sublinear integration in those cells regardless of stimulus congruency. It should be noted that especially in the OG condition, the non-preferred stimulus often elicited no response on its own, implying that the linear sum of component responses was equal to the response to the preferred monocular grating. For binocular cells, we found that linear integration approximated the binocular responses best for OG and CFS conditions, the two binocular conditions with binocular incongruency (OG: p = 0.104, CFS: p = 0.364). For binocular cell responses to the CG condition, supralinear integration was the best fit, with the slope significantly exceeding 1 (β_LS_ = 1.17 ± 0.08 [mean + SE], p = 0.047). Together with the findings from Fig. 2, this suggests that the binocular facilitation to congruent stimuli in binocular cells constitutes a mild boost over the linear response integration that occurs even for incongruent stimuli in these neurons.

Finally, to highlight the differences in binocular response integration between the three eye preference groups, we repeated the linear sum regression analysis but now pooled trials across all binocular conditions. This analysis produced a clear hierarchy of integration strength: the linear sum slope of binocular cells exceeded that of contra-preferring cells which in turn exceeded that of ipsi-preferring cells (binocular: β_LS_ = 1.16 ± 0.05, contra-preferring: β_LS_ = 0.82 ± 0.02, ipsi-preferring: β_LS_ = 0.51 ± 0.03, p < 0.001 for all three pairwise t-tests).

### Responses to ipsilateral stimuli are often poor predictors of binocular responses

For cells with a strong monocular preference, responses to monocular stimuli on the non-preferred eye are often weak or absent, which raises the question how well responses to individual monocular stimuli (instead of the linear sum of monocular stimulus responses) predict responses to binocular displays. To address this question, we repeated our regression analysis from Fig. 3 but now used the response to one of the two monocular stimuli (either contra- or ipsilaterally displayed) that make up a binocular display as a predictor for the response to that binocular display. We find that in the overall population, ipsilateral stimulus responses (ISR) were very poor predictors of binocular responses (Coefficient of determination r^2^_ISR_ for CG: 0.03 ± 0.02, OG: 0.00 ± 0.00, CFS: 0.01 ± 0.01; estimate ± SD; Fig 4A and B). In contra-preferring cells, the predictive performance of ipsilateral stimulus responses was similarly bad (r^2^_ISR_ for CG: 0.24 ± 0.06, OG: 0.02 ± 0.01, CFS: 0.27 ± 0.05) and always significantly worse than that of the contralateral stimulus responses (CSR; r^2^_CSR_ for CG: 0.60 ± 0.04, OG: 0.60 ± 0.05, CFS: 0.49 ± 0.05, p < 0.001 for all r^2^_ISR_ vs. r^2^_CSR_ comparisons, bootstrap hypothesis test, 1000 bootstrap samples, MCC for 12 tests; Fig. 4C). Surprisingly, even in ipsi-preferring cells, ipsilateral stimulus responses were at best mediocre predictors of binocular responses (r^2^_ISR_ for CG: 0.22 ± 0.09, OG: 0.13 ± 0.06, CFS: 0.10 ± 0.08) – likely because many ipsi-preferring cells have reduced responses in binocular displays. In fact, for ipsi-preferring cells in the CFS condition, contralateral stimulus responses (i.e. responses to a monocular contralateral Mondrian) were significantly better predictors of binocular responses than ipsilateral stimulus responses (i.e. responses to the preferred monocular ipsilateral grating; r^2^_CSR_: 0.69 ± 0.15, r^2^_ISR_: 0.10 ± 0.08, p < 0.001) - even though the cells of the ipsi-preferring group were selected to have strong responses to just such ipsilateral gratings (but not selected to respond to contralateral Mondrians). Only for binocular cells did the ipsilateral stimulus response predict binocular responses as well as the contralateral stimulus response (p > 0.75 for all three binocular conditions). We conclude that ipsilateral stimulus responses are poor predictors of responses to binocular stimuli throughout V1 and LM.

## Discussion

### Summary

In this study, we investigated mouse V1 and LM L2/3 neuronal responses to three types of binocular stimuli with varying congruency: congruent gratings, incongruent gratings and a flashing Mondrian pattern in binocular conflict with a grating. First, we found that neurons with a strong monocular preference tended to respond homogeneously to binocular stimuli regardless of congruency. Responses of cells with a preference for contralateral stimuli responded similarly to monocular and binocular stimuli while responses of cells with a preference for ipsilateral stimuli were suppressed equally across all binocular conditions. Binocular suppression in cells with a strong monocular preference also caused binocular responses to fall short of the linear sum of the responses to the corresponding monocular display such that the general population of cells exhibited sublinear integration. In contrast to the overall population trend, cells responsive to monocular stimuli on both eyes mostly exhibited linear response integration, or even supralinear integration in case of congruent stimulation. While many V1 and LM neurons show congruency-indifferent binocular response modulations, the subset of binocular cells notably does show sensitivity to the congruency of stimuli.

### Comparison to findings in primates

Some of the phenomena reported in this study, while new in the mouse, are similar to those observed in primates. For example, (Dougherty et al., 2019) report binocular modulations in seemingly monocular cells in macaque V1, and the same study also reports stronger interocular suppression for cells with a strong monocular preference. However, in macaques, the suppression of cells with a preference for the contra- or ipsilateral eye seems to be roughly equal (Zhang et al., 2024) while suppression in our data is more prevalent in ipsilateral-preferring cells. It should be noted that in macaques, the transient response (<150 ms after stimulus onset) to binocular stimuli is congruency-unspecific while congruency-specific response modulations emerge only later (Cox et al., 2019). It is possible that with calcium imaging, an early, congruency-unspecific response segment is mixed with a later congruency-specific response segment due to temporal blurring and that longer stimuli than those used in this study (1s) could help unmix these signals.

### Comparison to other mouse studies

Our findings of congruency-indifferent processing in V1 and LM diverge from those reported by an earlier study that also uses two-photon calcium imaging besides electrophysiology (Montgomery et al., 2026). Specifically, (Montgomery et al., 2026) report strong binocular facilitation in mouse V1 L2/3 pyramidal cells by *incongruent* stimuli while using similar (albeit static) grating stimuli as used in this study. While Montgomery et al. employed longer stimuli over multiple seconds, this facilitation effect already became apparent within the first second after stimulus onset. We do not replicate this finding, but note that our study was conducted using drifting grating stimuli at full contrast to maximize the number of responsive cells and only analyzed responses from conditions that contained each cell’s preferred grating. Such gratings alone evoke strong responses, which could mean that cells were already maximally excited such that facilitation effects as those reported by (Montgomery et al., 2026) may be have been obscured (although they were still apparent for binocular cells in the congruent condition).

Besides gratings, we also used a flashing Mondrian pattern as a monocular stimulus which is a stimulus type we have used before (Bassler et al., 2026). This stimulus is globally incongruent to the grating stimuli but may exhibit local congruence to a grating presented to the other eye, depending on the rectangle shown at a specific point in time and location in the visual field. It was originally designed to be strongly perceptually suppressive in a binocular display in humans (Tsuchiya & Koch, 2005), however, what effects it may have on mouse perception is unknown. Based on the findings of this study, the neuronal suppression effects reported in (Bassler et al., 2026) are not exclusive to binocular displays with Mondrian patterns but can also occur in binocular displays consisting only of gratings (as others have reported previously, e.g. (Longordo et al., 2013; Zhao et al., 2013)). In fact, in our population of grating-selective cells, the CFS condition did not seem to evoke particularly unique responses.

### Outlook

Several questions remain open regarding mouse binocular processing in early visual areas. What is the function of the strong ipsilateral signal suppression during binocular vision? Would ipsi-preferring cells only respond maximally in freely moving animals if the contralateral eye was occluded, or the binocular parts of the two eye inputs were somehow decorrelated (e.g. by eye movements)? Furthermore, it remains to be seen if stronger congruency-specific effects can be found in visual areas outside of V1 and LM such as RL, which may have a preference for binocular stimuli presented near to the eyes (La Chioma et al., 2019). Finally, although not addressed in this study, it would be interesting to determine how mice perceive binocular congruent and incongruent stimuli as reportable in a behavioral paradigm, and whether their binocular perception conform to, or differs from, perception integration and rivalry in primate vision.

## Acknowledgements

This project has received funding from the European Union’s Horizon 2020 Framework Programme for Research and Innovation under the Specific Grant Agreement No. 945539 (Human Brain Project SGA3) to C.P.

## Supplement

**Supplementary figure 1.**
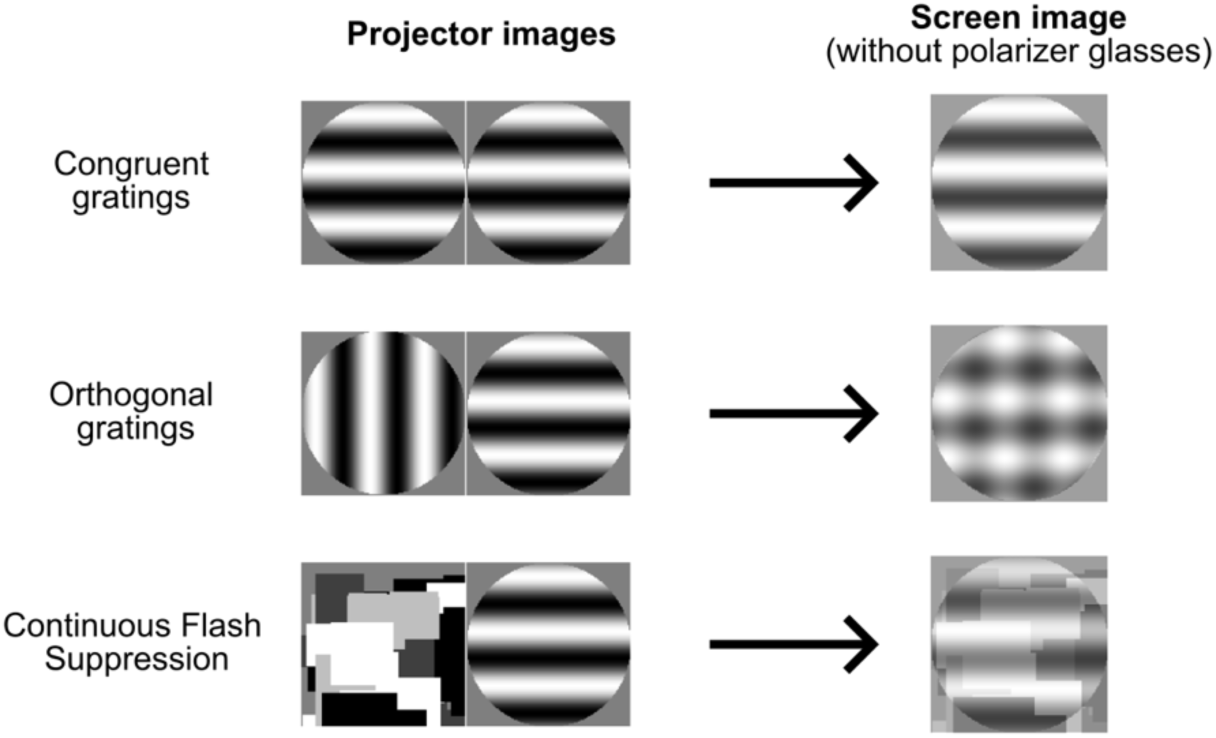
Illustration of projector images and screen images. To show binocular stimuli, two projectors were aimed at the same screen. Polarization filters were used to guide each projector image to one eye. Without polarization filter glasses, binocular stimuli appeared on the screen as an overlay of the two monocular components.

**Supplementary figure 2.**
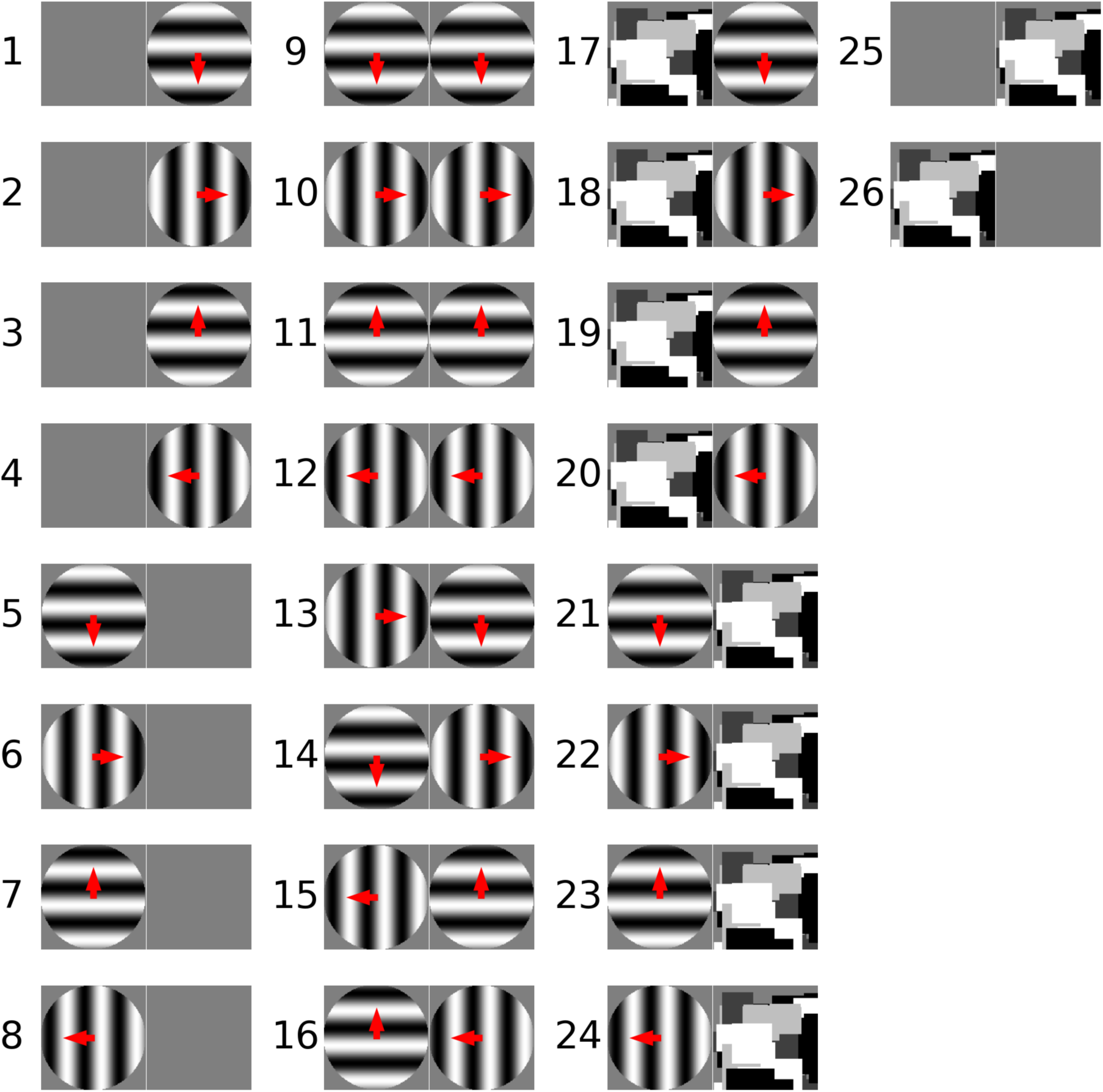
Overview over all stimulus conditions. The red arrow indicates the drift direction of the grating. Mondrian stimuli consisted of pseudorandomly arranged and sized rectangles with pseudorandomly assigned greyscale values. The rectangle arrangement was changed (“flashed”) every 100 ms. The sequence of rectangle patterns during the 1s of stimulus presentation was generated once at the beginning of very session, then kept constant.

**Supplementary figure 3.**
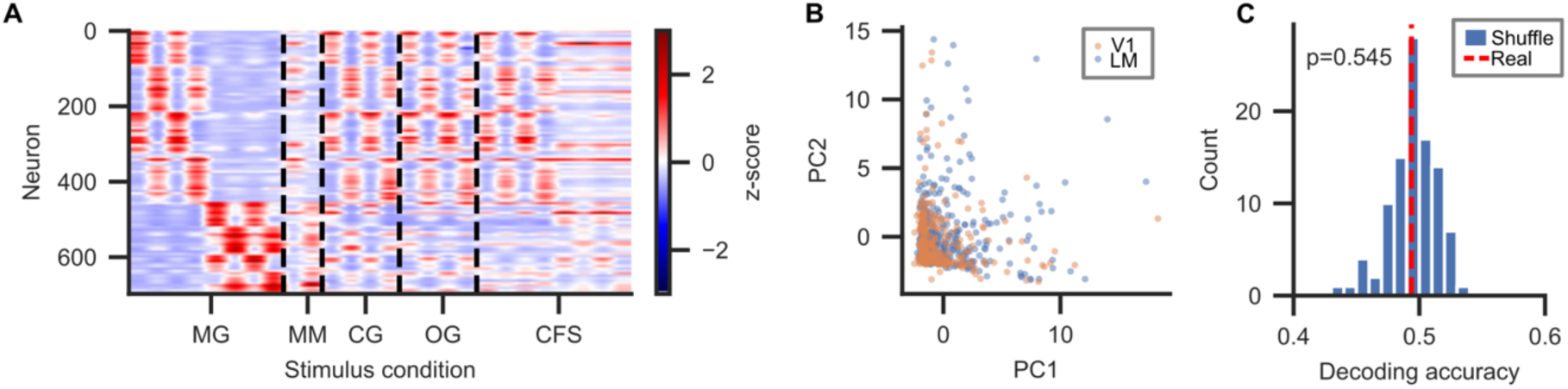
No decodable differences between V1 and LM cells. (A) Z-scored activation of all neurons to all 26 stimulus conditions. Cells were sorted by preferred grating for visualization purposes. (B) Scatterplot of principal component analysis (PCA) projections of (A) colored by brain area. Each dot represents a cell. V1 and LM dots do not form distinguishable clusters, suggesting that brain area is not a major source of variance in the data. PC: principal component. (C) Accuracy for decoding the brain area (V1 vs. LM) based on the average activity of each cell in each of the 26 stimulus conditions. The decoding accuracy was not different from chance level, indicating that V1 and LM cells could not be distinguished by their stimulus responses. Permutation test, 100 shuffles.

